# Biomathematical enzyme kinetics model of prebiotic autocatalytic RNA networks: degenerating parasite-specific hyperparasite catalysts confer parasite resistance and herald the birth of molecular immunity

**DOI:** 10.1101/2024.05.14.594066

**Authors:** Magnus Pirovino, Christian Iseli, Joseph A. Curran, Bernard Conrad

## Abstract

Catalysis and specifically autocatalysis are the quintessential building blocks of life. Yet, although autocatalytic networks are necessary, they are not sufficient for the emergence of life-like properties, such as self-propagation (replication) and adaptation. The ultimate and potentially fatal threat faced by molecular replicators is parasitism; if the polymerase error rate exceeds a critical threshold, even the fittest molecular species will disappear. Here we have developed an autocatalytic RNA early life mathematical network model based purely on enzyme kinetics, more specifically the steady-state approximation. We confirm previous models showing that these autocatalytic cycles are sustainable, provided there is a sufficient nucleotide pool. However, molecular parasites arise rapidly and become unsustainable unless they sequentially degenerate to hyperparasites (i.e. parasites of parasites). These hyperparasites acquire parasite binding specificity via two distinct temporal pathways. Our model is supported at three levels; firstly, ribozyme polymerases display Michaelis-Menten saturation kinetics and comply with the steady-state approximation. Secondly, ribozyme polymerases are capable of sustainable auto-amplification and of surmounting the fatal error threshold. Thirdly, with growing sequence divergence of host and parasite catalysts, the probability of self-binding increases and the trend towards cross-reactivity diminishes. Our model predicts that primordial host-RNA populations evolved via an arms race towards a host-parasite-hyperparasite catalyst trio that conferred parasite resistance within an RNA replicator niche. As such, it adds another mechanism – what’s more, with biochemical precision – by which parasitism can be tamed and offers an attractive explanation for the universal coexistence of catalyst trios within prokaryotes and the virosphere, heralding the birth of a primitive molecular immunity.

**Author Summary:** The quintessential components of life comprise a potent mixture of naturally occurring, but improbable chemical reactions (catalysis), and the arrangement of such accelerated chemical reactions into closed loops (autocatalytic sets). This is required, but is not sufficient for such networks to self-propagate (amplification of the information carrier = host polymerization) and adapt (Darwinian evolution). As soon as self-propagation is attained, the next hurdle is parasitism. This typically involves shorter molecules (the products of replicative errors) that hitchhike the replicative potential of the host. They will invariably outcompete the regular amplification process, unless a solution is found. We have addressed this problem using a new model based on the mathematics of catalysis. This model confirms previous studies demonstrating that autocatalytic sets become self-sustaining, assuming that a sufficient pool of molecular building blocks is available. However, molecular parasitism is pervasive and potentially fatal for both host and parasite. In our model, we allow these parasites to degenerate in a controlled fashion, giving rise to parasites of parasites (hyperparasites). As long as these hyperparasites acquire binding specificity for parasites, an attenuation of parasitism is observed. These parasite-hyperparasite cycles stabilize the host cycle, explaining why they are conserved, and why they are the likely reason behind the observation that all cellular hosts are associated with parasites (e.g. bacteria) and hyperparasites (e.g. viruses) across all kingdoms of life. Moreover, it provides a novel solution to the usually intractable problem of parasitism.

## Introduction

The multistep transition process from inanimate to living systems has been dated to >3.9 Ga [1]. A non-exhaustive list of thresholds that needed to be progressively surmounted during this process has been drawn, comprising, among others, chirality symmetry breaking, spontaneous polymerization, and self-assembly of compartments. The catalytic closure threshold and Eigen’s error threshold have received particular attention [2]. The latter posits that if the replication error exceeds a critical threshold value even the fittest replicators will not survive [3].

One central pillar in the theory of how early life could have materialized pertains to the “RNA world”. This stipulates that the initial informational polymer was RNA, and that the same molecule was also the fundamental catalyst [4]. The underlying concept of autocatalytic sets was laid out more than half a century ago [3,5]. It was subsequently refined [6] and validated in a proof of principle experiment showing that RNA fragments could build self-replicating ribozymes from individual RNA fragments via cooperative catalytic cycles; networks with three members showed strong, highly cooperative growth [7]. In parallel, an increasingly efficient number of RNA polymerase ribozymes were generated by *in vitro* evolution. These exhibited high levels of processivity [8], and displayed Michaelis-Menten saturation kinetics [9]. *In fine*, one RNA polymerase ribozyme surpassed Eigen’s critical error threshold, and successfully accomplished several rounds of cyclic self-replication [10,11]. Collectively, both theoretical considerations and experimental evidence support the view of a gradual evolution towards RNA self-catalysis. This started with self-assembly, followed by the formation of autocatalytic networks using self-assembled substrates. From this emerged different ribozymes (recombinases, ligases) and template based ligation, leading to a template based, self-replicase catalysed assembly of monomers [12].

The ribozyme catalysed RNA-polymerization process is even more error prone [4,8-11] than the proteinaceous RNA polymerases that notoriously lack proofreading activity [13]. This generates variant sequence space mutants called quasispecies [14,15]. This variant sequence-space intrinsically implies the obligate presence and evolutionary persistence of molecular parasites as observed in all life forms, and can be argued essentially for thermodynamic reasons [16,17]. Two main mechanisms have been proposed to confer parasite resistance to a given habitat, namely spatial self-organization [18-24], and physical compartmentalisation most probable via lipid bilayers, and the construction of protocells [25-27]. A direct comparison indicates that the survival of RNA replicators most likely occurred in protocells [28].

Since catalysis and autocatalysis are central to life, from a mathematical perspective the Michaelis-Menten reaction scheme appears pertinent [29], and implicitly the quasi steady-state assumption [30]. These considerations lay the ground for modelling catalysis, autocatalysis and the interactions within primordial, template-based ecological RNA communities using a single biomathematical tool derived from enzyme kinetics. We model here, based on a previously established tripartite population scheme [31], how molecular parasite resistance and a primordial molecular antiparasite immunity gradually emerged from progressively degenerating hyperparasite catalysts acquiring parasite-specificity.

## Results

Enzyme kinetics lies at the centre of the living world, and the intermediate complex lies at the centre of the centre. Consequently, in developing our current model we have used equations derived from enzyme kinetics, specifically the steady-state approximation by Briggs and Haldane [32]. Starting with a ribozyme polymerase exhibiting Michaelis-Menten saturation kinetics capable of beating the error-threshold [9-11], we demonstrate how template-based RNA autocatalytic cycles progressively degenerate to create successively parasite and hyperparasite cycles. Metabolic autocatalytic networks, arguably arising spontaneously and predating template-based polymerization [33], are assumed to be present and to provide sufficient energy and polymer precursors [23,31]. The previously established tripartite habitat framework, featuring a template-based RNA host and a molecular parasite that is itself parasitized (hyperparasite catalysts) [31] is instrumental for homeostasis and the emergence of molecular parasite resistance. It hinges upon hyperparasite catalysts acquiring parasite-specificity.

We provide mathematical models that describe the time evolution of autocatalytic RNA networks. The main goal of these models is to investigate the general conditions under which the stability of such networks could occur. The physical/chemical dimensions of the relevant input and output values (e.g. *t* = time, *k*_*cat*_, *k*_*on*_, *k*_*off*_, *K*_*M*_, RNA-concentration values, etc.) are assumed to be dimensionless. However, when available numerical values have been employed. Nevertheless, these values were not validated with respect to a specific experimental system.

### Biomathematical Michaelis-Menten kinetics and the Briggs-Haldane steady-state approximation for autocatalytic RNA networks

It is assumed that the ribozyme polymerase represents the positive RNA strand *R*_*1*_ with kinetic parameters capable of cyclic auto-amplification that surmount the error threshold (e.g. a *k*_*cat*_ of 0.1−1 min^−1^ and a *K*_*M*_ akin or superior to 0.1−1 μM) [9-11]. Its complementary negative RNA strand *R*_*2*_ is considered to be catalytically inert (uniquely a template). A complete autocatalytic replication cycle comprises the initial production of *R*_*2*_ by two *R*_*1*_ ribozyme molecules, and the subsequent reproduction of *R*_*1*_ by *R*_*1*_ using template *R*_*2*_ (Model 1, formalism described in Methods or Models, *R*_1_⇒ (*R*_1_*_R*_1_) →^*c*^ *R*_2_ *⇒* (*R*_1_*_R*_2_) →^*c*^ *R*_1_ (1.0), see also S1 Supporting Information for scheme in Fig 2A). Figure 1 shows a graphical representation of this model assuming realistic starting kinetics and habitat restriction parameters (described in Methods or Models). As can be seen, the abundance of these two molecular species increases rapidly and becomes stable, supporting previous work showing the viability of ribozyme polymerases, assuming that sufficient energy in the form of molecular building blocks is available [23].

**Figure 1.**
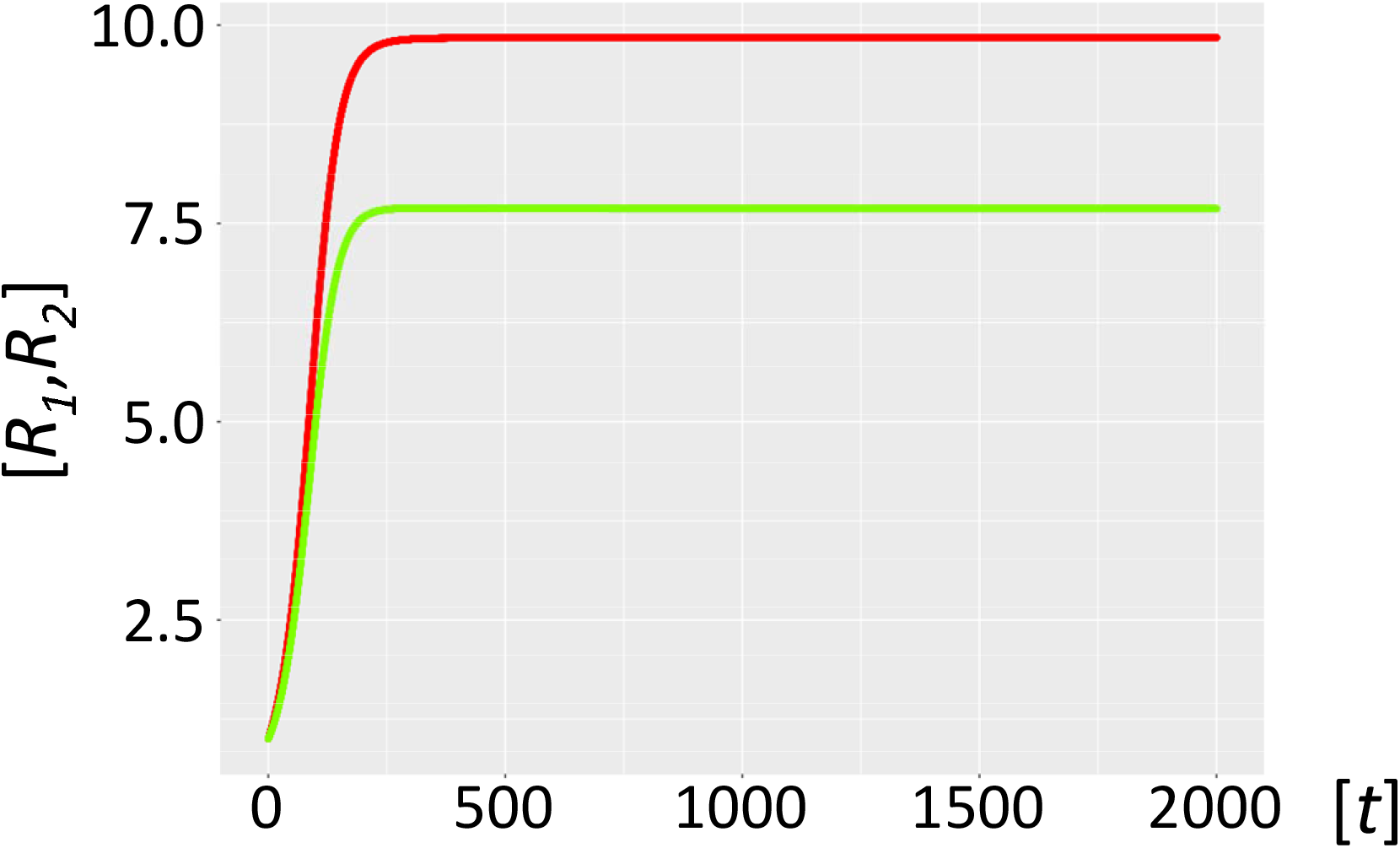
Habitat with autocatalytic host cycle (Model 1) Fig 1. Ribozyme concentrations of defined populations *R*_*1*_ and *R*_*2*_ [units on unspecified scale] are plotted against time *t* [units on unspecified scale]. The host catalysts (ribozyme polymerase *R*_*1*_, red, negative strand ribozyme *R*_*2*_, light green) build a self-amplifying and sustained autocatalytic habitat, as long as sufficient nucleotides are provided.

**Figure 2A.**
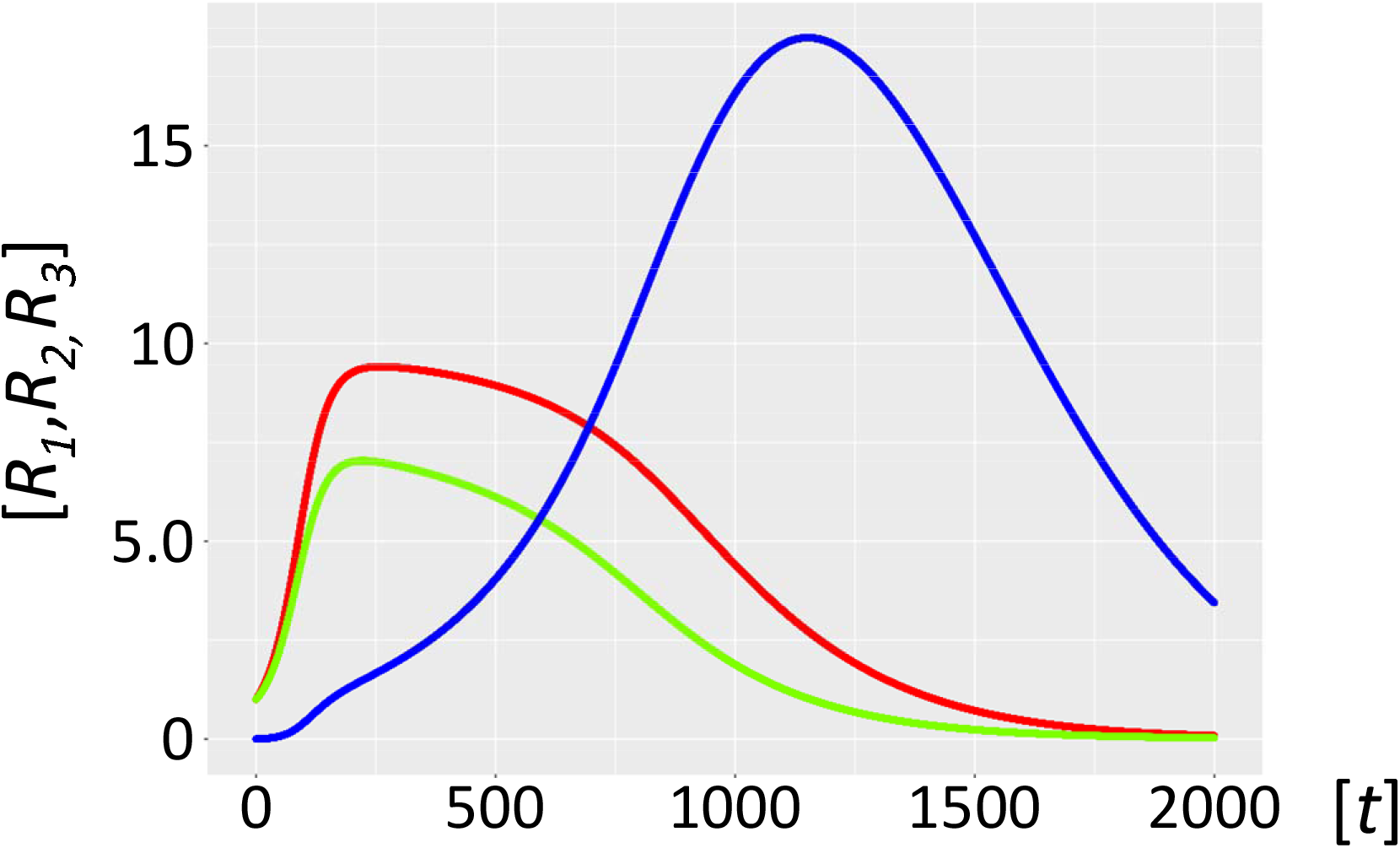
Autocatalytic habitat with molecular parasite exposure (Model 2.1) Fig 2A. Ribozyme concentrations of defined populations *R*_*1*_, *R*_*2*_ and *R*_*3*_ [units on unspecified scale] are plotted against time *t* [units on unspecified scale]. The host catalysts (ribozyme polymerase *R*_*1*_, red, negative strand ribozyme *R*_*2*_, light green) are rapidly invaded by the parasite ribozyme species *R*_*3*_ (blue), which is more efficiently amplified by the host ribozyme polymerase *R*_*1*_.

### Molecular parasitism in autocatalytic cycles

Molecular parasites are an inherent feature of the replication process. For a number of reasons they are inseparably associated with all life forms [16,17], with the most important being the accuracy-rate trade-off [34,35]. In general, enzymes operate orders of magnitude (*k*_*cat*_/*K*_*M*_ of *∼*10^5^ M^−1^ s^−1^ ) below the diffusion rate (10^8^ -10^9^ M^-1^ s^-1^ ). This implies promiscuous substrate binding, high catalytic activity, and as a secondary trade off, a high mutation rate generating a large variance in the quasispecies: i.e. speed being more important than enzyme accuracy [36]. Selection favours parasite molecules of subgenomic size [37-39], since the smaller they are the shorter their replication time, and the more overwhelmingly abundant they will become [38]. In addition, host ribozyme polymerases *R*_*1*_ amplifying parasites are unavailable for the synchronous host amplification cycle *R*_*1*__*R*_*2*_. This is exactly what Model 2.1 shows (Fig 2A, see also S1 Supporting Information for scheme in Fig 2B). It is merely a matter of time before the molecular parasites *R*_*3*_ = *P* outcompete the host molecules *R*_*1*_ and *R*_*2*_. Furthermore, if molecular parasites are diffusing in from a neighbouring habitat, the process is even more rapid (Model 2.2, Fig 2B). Therefore, as expected, high parasite loads overpower a typical habitat driven by efficient, but promiscuous ribozyme polymerases with *P* populations. This conundrum begs a solution.

**Figure 2B.**
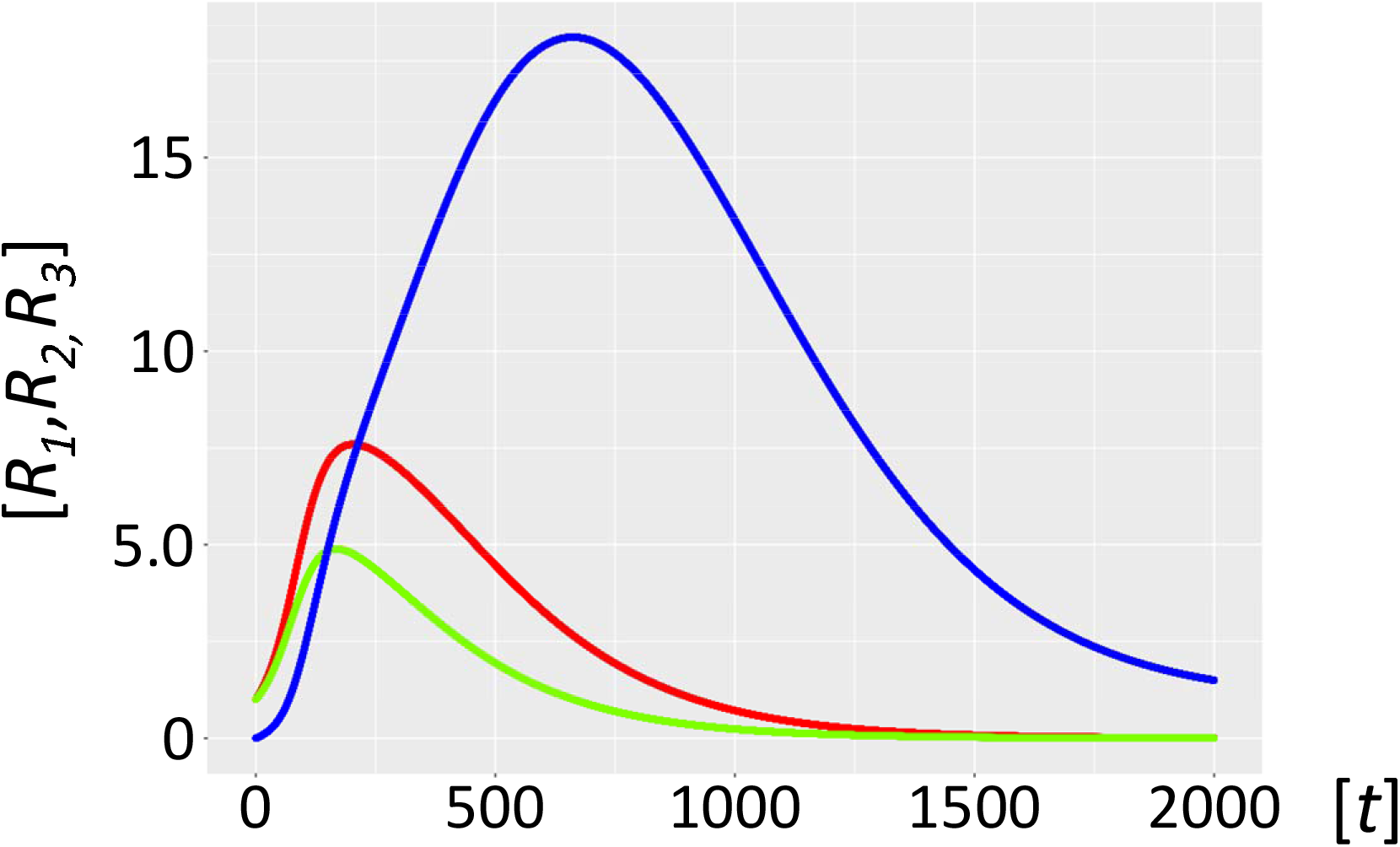
Autocatalytic habitat with molecular parasite inflow (Model 2.2) Fig 2B. Ribozyme concentrations of defined populations *R*_*1*_, *R*_*2*_ and *R*_*3*_ [units on unspecified scale] are plotted against time *t* [units on unspecified scale]. If parasite ribozymes *R*_*3*_ (blue) diffuse in from a neighbouring habitat, the host catalysts (ribozyme polymerase *R*_*1*_, red), and the negative strand ribozyme *R*_*2*_ (light green) are even more rapidly outcompeted and disappear.

### The emergence of hyperparasites: stable autocatalytic cycles in the presence of molecular parasites

The host ribozyme polymerase *R*_*1*_ is by design promiscuous ensuring amplification of both the host cycle *R*_*1*_*-R*_*2*_, and the very efficient amplification of the typically smaller *R*_*3*_ = *P* population (see above).Because of its high mutation rate, it will not only reproduce the original *P* species, but will inevitably also generate *P* mutants dubbed *F*, i.e. hyperparasites. Selection will preferentially amplify hyperparasite populations smaller than *P* [37-39] (Model 3, formally described in *Materials and Models, R*_1_ *→* (*R*_1__*P*) ⇒^*c*^*F*(1&2 ⇒ 3), see also S1 Supporting Information for scheme in Fig 2C). Short and loosely folded RNA sequences will be preferentially replicated (they are better templates), whereas longer and tightly folded ones may be better ribozymes and more resistant to hydrolytic decay [40]. As a corollary, *F* will be a highly efficient template, yet a poor or non-functional ribozyme capable of self-amplification. *R*_*1*_ is already efficiently producing *P* and *F*. However, this process is not subject to evolutionary pressure and is bound to degenerate (*F* ⇒ (*P*_*F*) *⇒*^*c*^ *nil F_F) →*^*c*^ *nil* (3.0)). The hyperparasite will parasitize *P* for several reasons; firstly, while bound in intermediate *P*_*F* complexes it is not available to *R*_*1*_, which can again engage in the amplification of the host cycle *R*_*1*_-*R*_*2*_. Secondly, in order for hyperparasitism to be highly efficient, it is required that *F* has a high specificity for *P*; i.e. in the presence of both *R*_*1*_ and *P* it binds mainly or exclusively to the latter. Indeed, this is precisely what the accuracy-rate trade-off predicts, with selectivity implying the ability to discriminate between two templates when both are present [35], whereas rate denotes speed [35,36]. Ground state discrimination means that specificity is achieved mainly through substrate binding, which imposes strong accuracy/rate trade-offs [34,35]. Improvements in selectivity mediated by tighter cognate substrate binding invariably leads to lower catalytic efficiency (parallel decreases in the constants for the cognate substrates *K*_*M*_^*cog*^ and *k* _*cot*_^*cog*^ [34,35]). This is precisely what the model stipulates; *F* has parasite- and hyperparasite-selectivity, which comes at the cost of reduced catalytic activity as would be predicted for a smaller molecular species. As shown in Fig. 3A (Model 3.1), and assuming the standard parameters as applied in Model 1, with similar *K*_*M*_ values (in the μM range) for both the intermediate complexes (*P_F* and *F_F*) and *R*_*1*__*R*_*2*_, hyperparasitism *per se* sustainably restores the abundance of the host cycle species *R*_*1*_ and *R*_*2*_. Still, the hyperparasite *F* is the most abundant species in the habitat, and the level of *P* remains important.

**Figure 3A.**
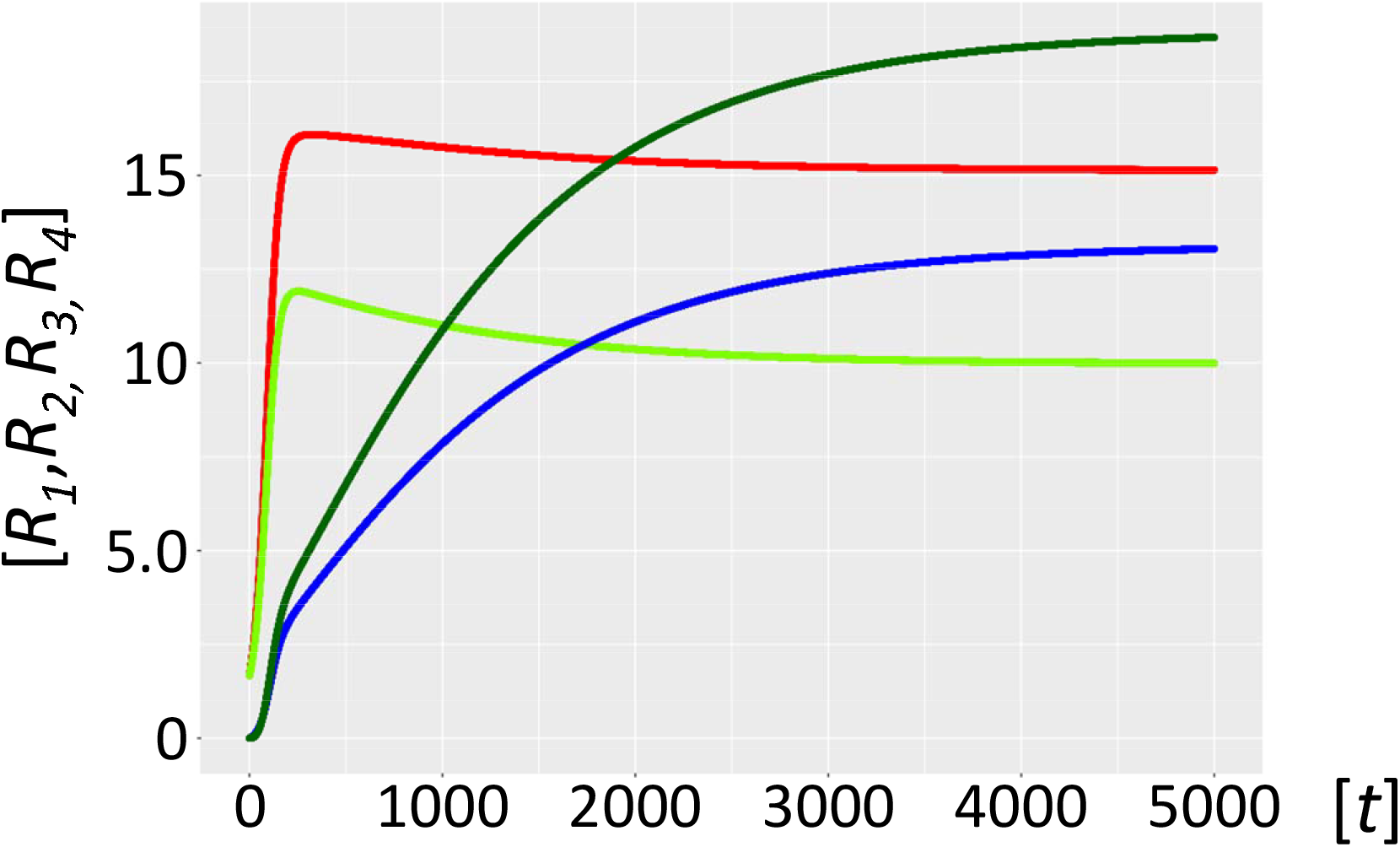
Stabilized host-parasite-hyperparasite cycle with *K*_*M*_ in μM range. Fig 3A. Ribozyme concentrations of defined populations *R*_*1*_, *R*_*2*_, *R*_*3*_ and *R*_*4*_ [units on unspecified scale] are plotted against time *t* [units on unspecified scale]. A habitat with highly efficient catalysts (ribozyme polymerase *R*_*1*_, red, negative strand ribozyme *R*_*2*_, light green) becomes stabilized by the generation and persistence of hyperparasites (parasite of parasite) *F* (dark-green) that parasitize the parasites *R*_*3*_ (blue). This already happens if intermediate complexes *P_F* and *F_F* have a *K*_*M*_ in the μM range (Model 3.1). However, the hyperparasite (dark-green) and parasite *R*_*3*_ (blue) ribozymes remain prevalent under these conditions.

However, increasing the binding affinity of *F* for itself and *P* by one order of magnitude (*K*_*M*_ values for *P_F* and *F_F* now in the 0.1 μM range, Model 3.2, Fig. 3B, see also S1 Supporting Information for scheme in Fig 2D) fully restores the abundance of the host cycle RNA species *R*_*1*_ and *R*_*2*_, at levels indistinguishable from the starting condition in Model 1. Since binding affinity under these conditions impacts directly on *K*_*D*_ values and hence catalytic efficiency (*k*_*cat*_/*K*_*M*_) [35], incremental changes in the affinity of *F* for itself and for *P*, by varying either the on-rate (fast association) or off-rate (slow dissociation) can impact significantly on the systems equilibrium [41,42]. Given what is currently known about template selection, modulating the on-rates appears to be the favoured option [43,44].

**Figure 3B.**
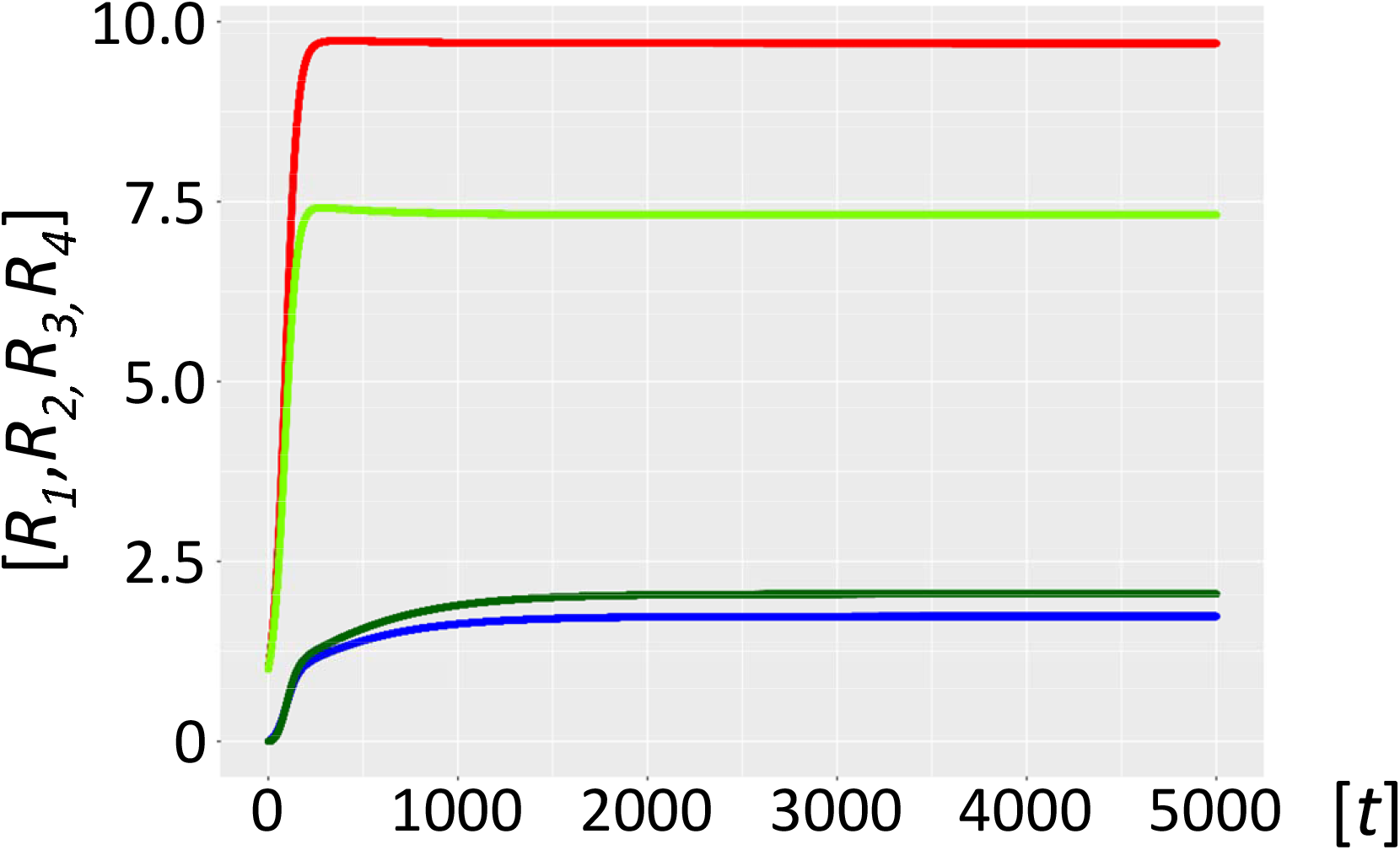
Stabilized host-parasite-hyperparasite cycle with *K*_*M*_ in 0.1 μM range. Fig 3B. Ribozyme concentrations of defined populations *R*_*1*_, *R*_*2*_, *R*_*3*_ and *R*_*4*_ [units on unspecified scale] are plotted against time *t* [units on unspecified scale]. A habitat with highly efficient catalysts (ribozyme polymerase *R*_*1*_, red, negative strand ribozyme *R*_*2*_, light green) becomes fully stabilized in the presence of parasite-hyperparasite cycles, if the *K*_*M*_ of the intermediate complexes *P_F* and *F_F* is decreased one order of magnitude in the 0.1 μM range (Model 3.2). Parasitism now is completely tamed (*P* blue, *F* dark green).

Several lines of experimental evidence suggest that self-binding within both host and parasite catalysts, with little cross-reactivity, is likely to happen. Firstly, just like proteins, RNAs exhibit a wide range of association rates, although even the fastest are significantly slower than those observed with proteins [42]. Nonetheless, RNA dimer formation is among the most rapid [42]. Secondly, the host and parasite-hyperparasite catalyst populations will diverge considerably, in both size and sequence, making the formation of homodimers or -multimers much more likely than that of heterodimers- or - multimers. Indeed, self-similarity increases the likelihood of physical interactions and spatially similar molecular structures interact preferentially [45]. In conclusion, with growing sequence divergence of host and parasite catalysts the probability of self-binding increases, and the trend towards cross-reactivity diminishes, precisely as suggested by our model.

Two temporally distinct scenarios can be envisaged; the parasite and hyperparasite immediately loose polymerase function because of the inherently high mutation load associated with the elevated replication rates of parasitic templates, and acquire parasite binding specificity in a second step. Consistent with this notion, the same laws that govern substrate-enzyme interactions are also valid for protein-protein associations [41,47], and as an extension here for ribozyme polymerases [34,43,44]. This scenario is discussed in in the main model. Alternatively, as outlined in the model described in S1 Supporting Information, hyperparasite catalysts with higher binding affinity for parasites are initially selected, but rapidly loose catalytic activity because of the accuracy-rate trade-off [35]. Subsequently, *F*’s catalytic activity is bound to degenerate, since *R*_*1*_ mediated replenishment of the parasite pool is far more efficient than the one resulting from the poor catalytic activity of *F*.

In sum, these simulations show that parasite resistance can be modelled with biochemical precision, and it relies on hyperparasites acquiring parasite binding specificity, via at least two temporally distinct pathways.

## Discussion

We present a novel early-life model for autocatalytic ribozyme polymerase networks based on enzyme kinetics, more specifically the steady-state approximation. As a starting point, we defined a ribozyme polymerase with kinetic parameters drawn from in vitro models, which was capable of sustaining cyclic self-amplification and overcoming the error-threshold. We subsequently confirmed earlier models showing that these autocatalytic cycles are sustainable, as long as a sufficient nucleotide pool is available. Inevitably, an efficient but promiscuous polymerase ribozyme with inherently low fidelity will generate molecular parasites that will overwhelm the habitat. However, because molecular parasitism is a chain process, parasites of parasites (hyperparasites) will eventually appear. As long as these hyperparasites acquire parasite binding-specificity, an increasingly solid and molecularly specific parasite resistance will ensue. We demonstrate that this parasite resistance can be modelled with biochemical precision. Collectively, these findings add one more mechanism by which parasites can be tamed. It offers an attractive explanation for the universal and tight presence of molecular trios within the cellular hosts from prokaryotes to the virosphere, and showcases the birth of parasite-resistance and a primitive molecular immunity (see below).

Once highly processive and efficient RNA catalysts are continuously provided to an experimental habitat, the host and parasite populations turn into an evolving ecosystem through Darwinian evolution, forming distinct host and parasite lineages that exhibit arms-race dynamics [37]. Resonating with this is the fact that self-replicating ribozyme networks consisting of three members display particularly high cooperative growth dynamics [7]. Intriguingly, there appears to be a trade-off between short and loosely folded sequences that are preferentially amplified (templates), and longer, tightly folded ones that are better ribozymes and are more resistant to hydrolytic decay [40]. This in-turn provides a rationale for splitting RNA communities into subgroups [7,37,40].

At the core of our current model is enzyme kinetics. The catalyst accuracy-rate trade-off analysis [35] applied to autocatalytic RNA networks [34] predicts the emergence of active ribozymes [10,11,46] with binding promiscuity and high catalytic activity, co-existing with subgenome-sized molecules characteristic of parasites with relatively lower catalytic activity [40]. This is precisely what our model predicts. Furthermore, this enzyme kinetics based model allows us to adapt these processes with high precision, since the input parameters can be varied at will. An incremental change in the affinity of the hyperparasite ribozyme for itself and for first generation parasite ribozymes can be achieved by varying association (fast association) and/or dissociation rates (slow dissociation) [41,42,47] although current models would favour the former option [43,44].

While primitive tripartite populations featuring rapidly degenerating catalysts likely represented a homeostatically stabilizing necessity, parasite communities within the population might have subsequently evolved more freely, with some becoming replicationally autonomous. Indeed, tripartite microbial populations composed of a host, a parasite and a hyperparasite are virtually ubiquitous [48], as evidenced for bacteria-bacteriophage-phage-satellites [49,50], host-RNA viruses-defective interfering RNA viruses [51], and eukaryotic host-NCLDVs-virophages [52]. Furthermore, it is striking that the appearance of microbial immunity is associated with hyperparasites. Several recent studies have shown that phages and their satellites encode a set of diverse antiphage systems [53,54], and defective interfering RNAs activate innate immunity [55]. Phage satellite-encoded immune systems protect bacteria from phage predation and were proposed to be an integral component of innate immunity [56]. This notion could easily be extended to omnipresent, bona fide hyperparasites, such as endogenous retroviruses [57,58]. Therefore, there is likely a strong, ancestral link between hyperparasites and parasite immunity.

We previously developed a tripartite early-life host-parasite-hyperparasite framework based on Lotka Volterra (LV)-equations [31,59]. The present work arrives at similar results, showing that LV-based equations provide an adequate approximation for the more realistic, kinetics-based model developed here. Nonetheless, the present model has by design intrinsic limitations, notably the fact that we restrict the reaction to one intermediate polymerase-template complex [60]. Additionally, we assume that the catalytic efficiency of ribozyme polymerases will slowly approach that of proteinaceous holoenzymes and of mainstream, albeit complexed, multi-subunit enzymatic machineries. These aspects will be addressed in future studies.

## Methods or Models

### Definitions and Terms

#### 1. Mathematical notation

If not alternatively specified, n-vectors are always column-vectors. If a real valued column-vector *xϵ*ℝ^*n*^ is transposed to a row-vector we write *x*^*T*^ . Accordingly, also *n*x*m* matrices *A ϵ* ℝ^*nxm*^ can be transposed by the same symbol *A*^*T*^ . The *i*_*th*_ entry of a vector is denoted by *x*_*i*_. The entry of the *i*_*th*_ row and the *j*_*th*_ column of a matrix *A* is denoted by *A*_*ij*_ .

#### 2. Michaelis-Menten and Briggs-Haldane kinetics

The enzyme kinetics generally known as Michaelis-Menten kinetics [29,30] is introduced as follows:

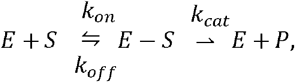

*E* being the enzyme, *S* the substrate, *E*_*S* the intermediate enzyme-substrate complex, and *P* the product. If all the symbols here are meant to be concentrations in a given environment (habitat), then the following equations apply:

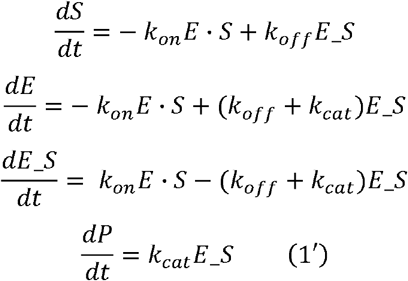

An important assumption is that the concentration of *E*−*S* reaches the steady-state almost instantly (steady-state approximation). According to the Briggs-Haldane steady-state approximation [32], we therefore assume

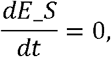

and thus:

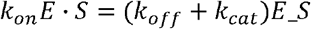

Here *E* is the free enzyme *E*, and *S* the free substrate. If we write

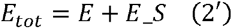

then the equation above yields, with the Michaelis constant 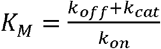

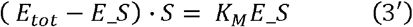

and thus

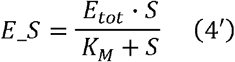

#### 3. Michaelis-Menten kinetics in the RNA-world

Let us now consider a set of different RNA populations { *R*_1,_ *R*_2,_ *…, R*_*n*_ } in a primordial molecular RNA landscape, with the supply of nucleotides being provided by metabolism [23,31]. This metabolism is not further specified, and we assume that there is a limitation on the number of RNA molecules { *R*_1,_ *R*_2,_ *…, R*_*n*_ } that can be maintained (habitat restriction constant, see below). These RNA molecules may be ribozyme polymerases if they exert a catalytic activity or merely templates.

##### 3.1 A simple three ribozyme system

In this RNA world, a catalytic activity is exerted if a free ribozyme polymerase exhibiting Michaelis-Menten saturation kinetics [9] from population *R*_1_ encounters another from *R*_2_. The ribozyme *R*_1_ uses the free molecule *R*_2_ as template, and by forming a polymerase-template complex, catalyses the production of a third ribozyme population *R*_3_ (*R*_3_ being the negative strand of *R*_2_) We now apply Michaelis-Menten kinetics, which entails that not only *R*_1_ is a ribozyme polymerase, but also the substrate *R*_2_, and even the product *R*_3_. The Michaelis-Menten formalism is valid for this special example. If *R*_1_, *R*_2_, *R*_3_ denote the concentrations of the total ribozyme masses, and 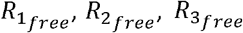 denote the concentrations of their respective free ribozyme masses, then 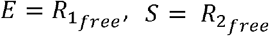, and *P* =*R*_3_. Then, equation (2’) reads as follows: 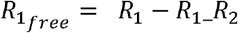 . But now, the free substrate 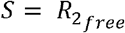 is also a function of the total substrate *R*_2_ and its enzyme-substrate complex *R*_1__*R*_2_, i.e.,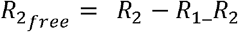 . Equations (1’) and (3’) from above now read as follows:

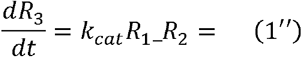

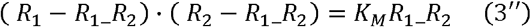

According to Briggs-Haldane it is assumed that the enzyme-substrate complex *R*_1__*R*_2_ reaches the steady-state very rapidly, 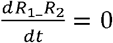 Equation (3’’) is now a nonlinear quadratic equation, but still solvable for the unknown *R*_1__*R*_2_:

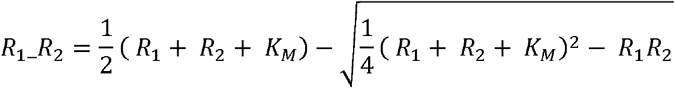

This solution of equation (3’’) is called the tight-binding or Morrison equation [61].

##### 3.2 The general case of n ribozymes

In the above special case of three ribozymes, we have assumed that only ribozymes *R*_1_ and *R*_2_ are forming an enzyme-substrate complex *R*_1__*R*_2_. Ribozyme *R*_1_ does not bind to ribozyme *R*_3_ or to itself *R*_1,_nor does ribozyme *R*_2_ bind to *R*_1_ or to *R*_3_. In the general case, where we have *n* ribozymes *R*_1,_ *R*_2,_*… R*_*n*_, every ribozyme polymerase *R*_*i*_ could potentially bind - through its role as enzyme - to a template (substrate) ribozyme *R*_*j*_, forming an enzyme-template complex *R*_*i_*_*R*_*j*_. In general, complex *R*_*i_*_*R*_*j*_ is not equal to complex, *R*_*j_*_*R*_*i*_ since *R*_*i_*_*R*_*j*_ is governed by the enzymatic activity of *R*_*i*_, whereas *R*_*j_*_*R*_*i*_ may be governed by the different enzymatic activity of *R*_*j*_, which may not be the same.

Thus, in general, *n* ribozymes can bind to *n*^2^ different enzyme-template complexes *R*_*i_*_*R*_*j*,_ *i,j* =1, …,*n*.

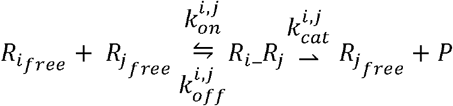

Equation (1’), describing the production rate of new products, now becomes

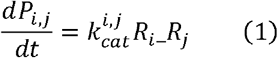

In an *n* ribozymes RNA world, the products *P*_*i,j*_ can be found among the given ribozymes, i.e., *P*_*i,j*_ ∈ {*R*_1_,*R*_2_, …,*R*_*n*_} . Equation (2’), describing the relation between the total (*R*_*i*_) and the free 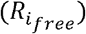 concentration of ribozymes, now turns into the following equation(2) :

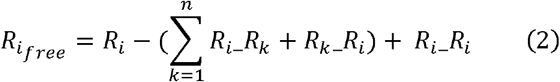

Again, we assume according to Briggs-Haldane that all ribozyme-template complexes *R*_*i_*_*R*_*j*_ reach the steady-state very rapidly, 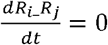 *i,j = 1, …, n* .

Given the Michaelis constants 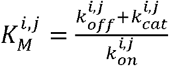, equation (3’) turns now, for every *i,j* = 1, …, *n*, into

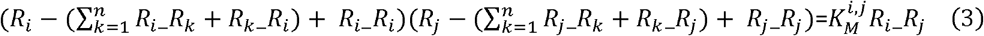

Equation (3) is a nonlinear system of *n*^2^ equations with *n*^2^ unknown ribozyme-template complexes *R*_*i_*_*R*_*j*_ Given the total concentrations, *R*_1,_ *R*_2,_ *… R*_*n*_, this equation is solvable through an appropriate nonlinear multidimensional multistep algorithm. For this purpose, we propose a Newton iteration outlined as follows.

##### 3.3 Newton iteration for the solution of equation (3)

Let 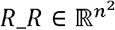 be a real valued *n*^2^ dimensional vector, with

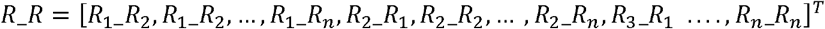

Let 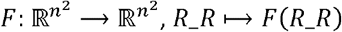, be a *n*^2^-multidimensional function 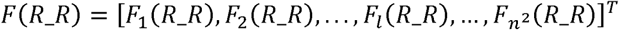 with

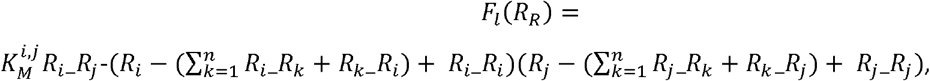

*l*=1,…, *n*^2^, where the indices *i,j* themselves are functions of index *l*. If we define these functions in the following way:

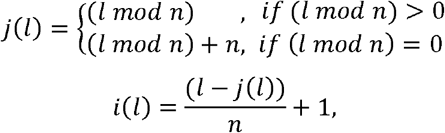

then the *l*th component of vector *R*_*R* is well-defined as (*R*_*R*)*l* = *R*_*i*(*l*)__*R*_*j*(*l*)_, and also the *l*th component of *F,F*_*l*_(*R*_*R*) is well defined for every index *l*=1, …,*n*^2^ . Now we state that a vector *R*_*R* = [*R*_1_ _ *R*_2_, *R*_1_ _ *R*_2_, … *R*_*n_*_*R*_*n*_]^*T*^ is a solution of (3), if an only if *F* (*R*_*R)=* 0,*i*.*e. F*_*l*_*(R*_*R)*= 0 for every *l*=1, …,*n*^2^. The classic multidimensional Newton iteration algorithm needs an explicit expression for of the Jacobian *J* of *F(R*_*R):*

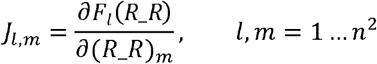

The full derivation of the formula for the Jacobian is straight forward, therefore we just give here the final result. Let 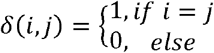be the Kronecker symbol, then the explicit formula for the Jacobian is given as follows. For *l,m* = 1, …,*n*^2^:

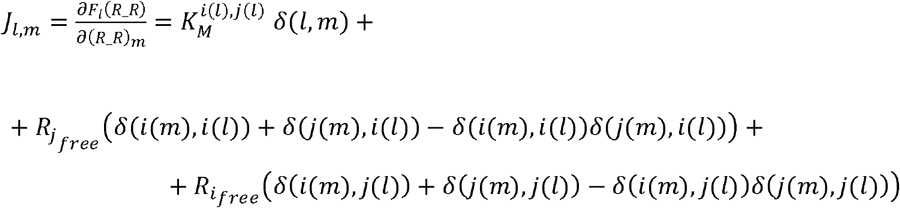

In this formula, the free ribozymes, 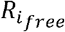, are calculated according to (2).

Now we are able to write down the full Newton iteration algorithm. Given some initial set of total ribozyme concentrations { *R*_1_, *R*_2_,… *R*_*n*_}, we start assuming that no ribozyme-template complex has been built yet; i.e. 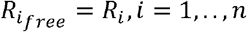, and (*R*_ *R)*_*l*_ = 0, *l* = 1,…,*n*^2^ The initial vector of ribozyme-template complexes therefore equals to zero: (*R*_ *R)*^(0)^ = 0. After *k* iterations, the resulting vector of ribozyme-template complexes is then calculated by:

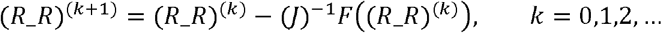

Here (*J*)^−1^ is the is *n*^2^ × *n*^2^ inverse matrix of the Jacobian *J*. Normally, this sequence of iterations convergences to a solution (*R*_ *R)*^*(k )*^ → (*R*_ *R)*^*\**^ =*R*_ *R* with *F*((*R*_ *R)*^*\**^ ) = 0.The entries of the resulting vector *R*_*R* are then concentrations of all of the possible ribozyme-template complexes (*R*_ *R)*_*l*_ = *R*_*i(l)*__*R*_*j(l)*_ *1,…,n*^2^ solving equation (3).

## 4. Template based autocatalytic cycles

### 4.1 Preamble

Reconsider the set of populations of different RNA molecules { *R*_1,_ *R*_2,…_ *R*_*n*_}, introduced above. The ribozymes form a catalytic cycle, if

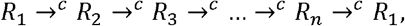

where the symbol “*A*→^*c*^ *C*” means: ribozyme *A* catalyzes ribozyme *C*. The catalytic production of *C* is called template based if an encounter with a template ribozyme *B* is needed for this process, i.e. *A* needs to form, together with *B*, an enzyme-template intermediate complex *A_ B* in order to produce *C*. We write “*A_B*→^*c*^ *C*” . If the autocatalytic cycle reproduces not only all ribozymes carrying an enzymatic role, but also all ribozymes serving as templates along its way then we call it a template based autocatalytic cycle. The following example meets all the criteria of such a template based autocatalytic cycle:

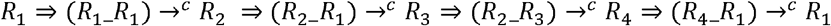

This reads as follows, a ribozyme *R*_1_ forms an enzyme-template complex with itself, *R*_1__*R*_1_, producing a new copy of ribozyme *R*_2_ . *R*_2_ too forms an enzyme-template complex with *R*_1_, *R*_2__ *R*_1_, producing ribozyme *R*_3_. *R*_2_ now forms another enzyme-template complex with *R*_3_, *R*_2_ _ *R*_3_, producing *R*_4_, which in turn produces again *R*_1_ by forming a catalytic enzyme-template complex *R*_4__ *R*_1_ with *R*_1_ . Now we are able to model primitive template based autocatalytic cycles in an RNA landscape, with a given, but not further specified metabolism providing a limited supply of nucleotides.

### 4.2 Model 1: the host cycle

Let ribozyme *R*_1_ be a population of positive strand ribozyme polymerases, and *R*_2_ be the population of its negative strand, then we can model the following template based autocatalytic cycle:

**Model 1**

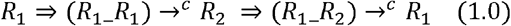

The replication dynamics is governed by the following ODEs:

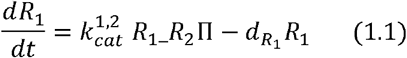

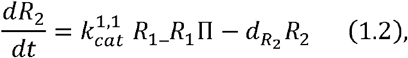

Here 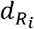 are the decay rates of populations *R*_*i*_,*i =*1,2, and 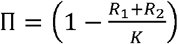 is the habitat restriction constant. *K*is the maximum number of *R*_1_ or *R*_2_ RNA molecules the habitat is able to maintain through the given metabolism. Given the total concentrations *R*_1_ and *R*_2_ we assume using Briggs-Haldane that the enzyme-template intermediate complexes *R*_1__ *R*_1_ and *R*_2_ *_R*_2_ are reaching their steady-state at every point in time very rapidly. In other words, we assume that the process of attaining the steady-state for the enzyme-template intermediate complexes is by some orders of magnitudes faster than the production of new ribozymes. In order to calculate all of the possible enzyme-template complexes *R*_*i*_ *_R*_*j*_ through the Newton iteration algorithm, as outlined above, we need to attribute values to 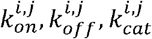, along the Michaelis constants 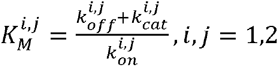. E.g.

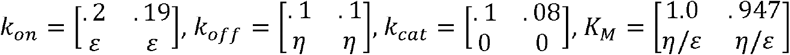

We further note that in this model ribozyme *R*_2_ does not form an enzyme-template complex with *R*_1_, nor with itself, assuming that *R*_2_ - the negative strand of *R*_1_ - does not have catalytic activity. Since the Newton algorithm needs Michaelis constants for all possible enzyme-template complexes *R*_*i*__*R*_*j*_, we assume here that 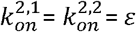 is a very low number greater than zero, and also 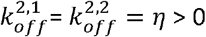. We choose *ε* and *η* so that formation of enzyme-template complexes *R*_2__ *R*_1_, *R*_2__ *R*_2_ is possible, but always at very low concentration and also dissociating very rapidly; e.g., *ε* = 10^−5^, *η*= 10^−1^. By choosing appropriate values for all other parameters (e.g. 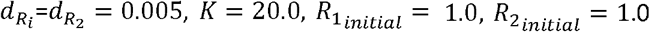 ). The ODEs (1.1), (1.2) of Model 1 are solved numerically by simple timestep integration. Note that for every timestep *t*_*k*_=*t*_0_ + *k* Δ*t* a calculation of all the possible enzyme-template complexes *R*_*i*_ *_R*_*j*_ through the Newton iteration algorithm is needed.

### 4.3 Model 2: the emergenc of parasitism

Autocatalytic host cycles (1.0) inevitably are subject to molecular parasitism [16,17]. In (1.0) the ribozyme polymerase *R*_1_ is replicating *R*_2_ and itself. Such a replicase cannot be so specific that it only replicates *R*_1_ and *R*_2_, since under such specific conditions *R*_1_ would quickly lose its catalytic activity due to mutations it is subject to [16,17]. Therefore, ribozyme polymerase *R*_2_ will inevitably be parasitized by RNA templates, typically of subgenomic size [37,38], say *R*_3_ = *P* (parasites):

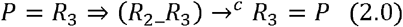

This motivates following additional equation

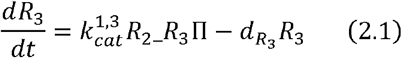

The habitat restriction term Πextends to

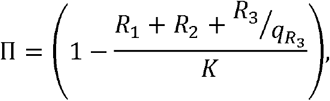

wher 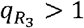, since parasites *R*_3_ = *P* are more efficient at using the habitat resources, because of their smaller size that confers a replicative advantage [38]. A ribozyme polymerase *R*_1_ that engages in replicating *R*_3_ =*P*, building an enzyme-template complex *R*_1__ *R*_3_, cannot simultaneously form enzyme-template complexes with *R*_1__ *R*_2_ or *R*_1__ *R*_2_. Thus, *R*_3_ parasitizes the whole autocatalytic cycle (1.0) . Here, for simplicity, we do not distinguish between positive and negative strand parasites.

We model two parasite sources, the most probable being the imperfect fidelity of polymerase *R*_1_. Rather than producing a viable RNA molecule *R*_2_ or *R*_1_, it may, at a specific mutation rate, *μ*> 0, produce a non-viable mutant having more similarity to a parasite *P* =*R*_3_, rather than to a viable RNA molecule of the host cycle (1.0). Parasite invasion from a neighbouring habitat can be viewed as an alternative.

The ODE equations for Model 2 are as follows:

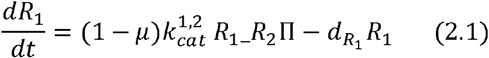

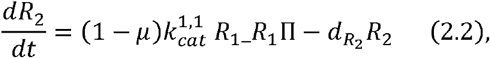

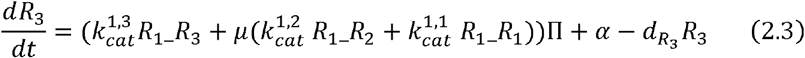

Here we assign values corresponding to the ones used in Model 1: 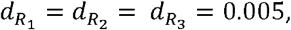 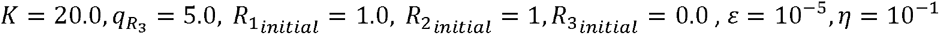, and

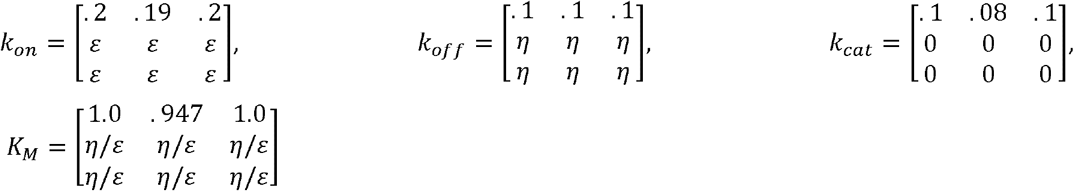

### 4.4 Model 3: the emergence of hyperparasitism

Parasitism is a chain process, molecular parasites of even smaller size than the first generation parasites are bound to emerge over time [37,38]. The model stipulates that the ribozyme polymerase *R*_1_ replicates host templates *R*_1_ faithfully the enzyme-template complexes *R*_1_*_R*_1_ and *R*_1_*_R*_2_ only catalyze *R*_2_ and *R*_1_, respectively. If replicase *R*_1_ binds to a parasite template *R*_1_*_R*_3_, it not only produces another copy of *R*_3_, or a negative strand of *R*_3_, respectively, which we again for simplicity do not distinguish here, and also another, smaller RNA molecule *R*_4_ =*F*, which we call here the hyperparasite, *F* being smaller than *R*_3_, but partially overlapping with it.

The hyperparasite cycle is triggered from the following process

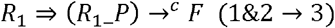

and has the following form:

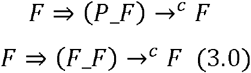

The hyperparasite *F* binds to an enzyme-template complex *P_F* or to itself *F_F*, which implies that *F* is capable of replicating itself. Again for simplicity, we do not distinguish between positive and negative strand hyperparasites *R*_4_ =*F*, assuming both are present. But since the production of *F* is already guaranteed through the trigger process(1&2 →3), such a process may degenerate to produce futile intermediate complexes [62] *P_F* or *F_F*:

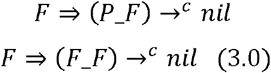

*P* = *R*_4_ is a hyperparasite, because it parasitizes the parasite *P* while bound to a complex *P_F*. A parasite molecule *P* bound to *F* is not able at the same time to bind to the ribozyme polymerase *R*_1_, and therefore cannot be replicated anymore. In order to be effective for the host, the hyperparasite requires to have binding specificity for parasite molecules, *R*_3_ = *P: F* : should form intermediate enzyme-template complexes only with *R*_3_ = *P* molecules or with itself.

Now we are able to write down the ODE equations for Model 3, the host-parasite-hyperparasite model.

**Model 3**

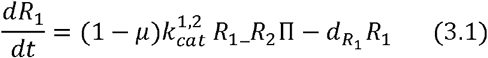

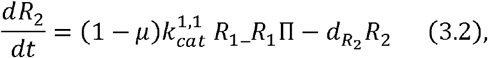

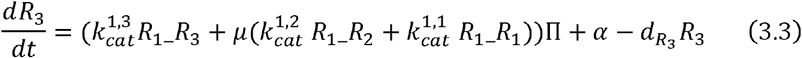

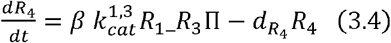

The habitat restriction parameter extends here to

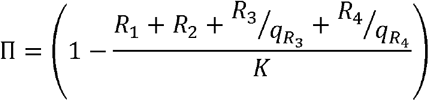

Here we again assign values accordingly to the ones in Model 1 and 2: 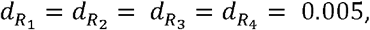 with 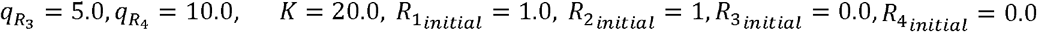, mutation rate *μ* = .01, parasite inflow per unit of time, *α* = 0.001, *β* =1.5 (*β* is here a parameter quantizing the trigger process (1&2 → 3) ), *ε* = 10^−5^, *η* = 10^−1^ and

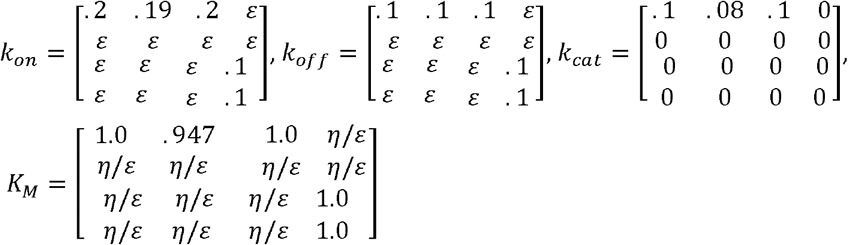

The values for Model 3 are exactly the same as in Model 2, except for the parameters affecting the hyperparasite *R*_4_ =*F* .

### 4.5 Model 3.1, μM-range Michaelis constants for P_F and F_F complexes

Here we chose the following parameters for the binding of the hyperparasite *R*_4_ =*F* with *R*_3_ = *P* or with itself: 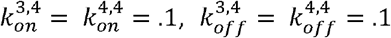 and hence the Michaelis constants 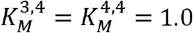 .

### 4.6 Model 3.2, sub-μM-range Michaelis constants for P_F and F_F complexes

Now if the binding of the complexes *P_F, F_F* is further strengthened, by e.g. increasing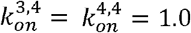, and so lowering the Michelis constants 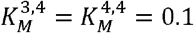, the situation for the host cycle can even be further stabilized, with a lower population of parasites *P* and less molecules bound to the *P_F* and *F_F* complexes. Thus, if the hyperparasite *F* does not or only very weakly interact with the larger host ribozyme polymerases *R*_1_ and *R*_2_ but does bind with higher affinity to parasite molecules *P* and also to itself, then it is offering solid stability to the host cycle in the presence of parasites *P*.

## Acknowledgements

We thank Thomas Curran for discussions.

## Author contributions

Conception and design of the work: MP and BC. Data acquisition MP; data analysis MP, BC, JAC, CI; interpretation of data MP, BC, JAC, CI. Creation of new software used in the work: MP, CI. Drafted the work and revised it: BC, JAC and MP.

## Data availability

The data, including scripts, is fully made available at the following sites (https://github.com/BICC-UNIL-EPFL/biomathematical-enzyme-kinetics-model [github.com], and the DOI: https://doi.org/10.5281/zenodo.11060828 [doi.org].

## References

1. Betts HC, Puttick MN, Clark JW, Williams TA, Donoghue PCJ, Pisani D. Integrated genomic and fossil evidence illuminates life’s early evolution and eukaryote origin. Nat Ecol Evol. 2018;2: 1556–1562. Doi: 10.1038/s41559-018-0644-x. PMID:30127539

2. Jeancolas C, Malaterre C, Nghe P. Thresholds in Origin of Life Scenarios. iScience. 2020;23: 101756. Doi: 10.1016/j.isci.2020.101756. PMID:33241201

3. Eigen M. Self-organization of matter and the evolution of biological macromolecules. Naturwissenschaften. 1971;58: 465–523. PMID:4942363

4. Joyce GF, Szostak JW. Protocells and RNA Self-Replication. Cold Spring Harb Perspect Biol. 2018;10: a034801. Doi: 10.1101/cshperspect.a034801. PMID:30181195

5. Kauffman SA. Cellular homeostasis, epigenesis and replication in randomly aggregated macromolecular systems. J Cybernet. 1971;1: 71–96.

6. Kauffman SA. Autocatalytic sets of proteins. J Theor Biol. 1986;119: 1–24. Doi: 10.1016/s0022-5193(86)80047-9. PMID:3713221

7. Vaidya N, Manapat ML, Chen IA, Xulvi-Brunet R, Hayden EJ, Lehman N. Spontaneous network formation among cooperative RNA replicators. Nature. 2012;491: 72–7. Doi: 10.1038/nature11549. PMID:23075853

8. Tjhung KF, Shokhirev MN, Horning DP, Joyce GF. An RNA polymerase ribozyme that synthesizes its own ancestor. Proc Natl Acad Sci U S A. 2020;117: 2906–2913. Doi: 10.1073/pnas.1914282117. PMID:31988127

9. Kakoti A, Joyce GF. RNA Polymerase Ribozyme That Recognizes the Template-Primer Complex through Tertiary Interactions. Biochemistry. 2023;62: 1916–1928. doi: 10.1021/acs.biochem.3c00091. PMID:37256719

10. Papastavrou N, Horning DP, Joyce GF. RNA-catalyzed evolution of catalytic RNA. Proc Natl Acad Sci U S A. 2024;121: e2321592121. doi: 10.1073/pnas.2321592121. PMID:38437533

11. Chen IA. RNA life on the edge of catastrophe. Proc Natl Acad Sci U S A. 2024;121: e2402649121. doi: 10.1073/pnas.2402649121. PMID:38478681

12. Pavlinova P, Lambert CN, Malaterre C, Nghe P. Abiogenesis through gradual evolution of autocatalysis into template-based replication. FEBS Lett. 2023;597: 344–379. Doi: 10.1002/1873-3468.14507. PMID:36203246

13. Steinhauer DA, Domingo E, Holland JJ. Lack of evidence for proofreading mechanisms associated with an RNA virus polymerase. Gene. 1992;122: 281–8. doi: 10.1016/0378-1119(92)90216-c. PMID:1336756

14. Domingo E, García-Crespo C, Perales C. Historical Perspective on the Discovery of the Quasispecies Concept. Annu Rev Virol. 2021; 8: 51–72. Doi: 10.1146/annurev-virology-091919-105900. PMID:34586874

15. Bull JJ, Meyers LA, Lachmann M. Quasispecies made simple. PloS Comput Biol. 2005;1: e61. Doi: 10.1371/journal.pcbi.0010061. PMID:16322763.

16. Koonin EV, Wolf YI, Katsnelson MI. Inevitability of the emergence and persistence of genetic parasites caused by evolutionary instability of parasite-free states. Biol Direct. 2017;12: 31. Doi: 10.1186/s13062-017-0202-5. PMID:29202832

17. Iranzo J, Puigbò P, Lobkovsky AE, Wolf YI, Koonin EV. Inevitability of Genetic Parasites. Genome Biol Evol. 2016;8: 2856–2869. doi: 10.1093/gbe/evw193. PMID:27503291

18. Szabó P, Scheuring I, Czárán T, Szathmáry E. In silico simulations reveal that replicators with limited dispersal evolve towards higher efficiency and fidelity. Nature. 2002;420: 340–3. doi: 10.1038/nature01187. PMID:12447445

19. Könnyu B, Czárán T, Szathmáry E. Prebiotic replicase evolution in a surface-bound metabolic system: parasites as a source of adaptive evolution. BMC Evol Biol. 2008;8: 267. doi: 10.1186/1471-2148-8-267. PMID:18826645

20. Branciamore S, Gallori E, Szathmáry E, Czárán T. The origin of life: chemical evolution of a metabolic system in a mineral honeycomb? J Mol Evol. 2009;69: 458–69. doi: 10.1007/s00239-009-9278-6. PMID:19806387

21. Takeuchi N, Hogeweg P. Multilevel selection in models of prebiotic evolution II: a direct comparison of compartmentalization and spatial self-organization. PLoS Comput Biol. 2009;5: e1000542. doi: 10.1371/journal.pcbi.1000542. PMID:19834556

22. Shay JA, Huynh C, Higgs PG. The origin and spread of a cooperative replicase in a prebiotic chemical system. J Theor Biol. 2015 Jan 7;364:249–59. doi: 10.1016/j.jtbi.2014.09.019. PMID: 25245369.

23. Kim YE, Higgs PG. Co-operation between Polymerases and Nucleotide Synthetases in the RNA World. PLoS Comput Biol. 2016;12: e1005161. doi: 10.1371/journal.pcbi.1005161. PMID:27820829

24. Tupper AS, Higgs PG. Error thresholds for RNA replication in the presence of both point mutations and premature termination errors. J Theor Biol. 2017;428: 34–42. doi: 10.1016/j.jtbi.2017.05.037. PMID:28606750

25. Zintzaras E, Santos M, Szathmáry E. Selfishness versus functional cooperation in a stochastic protocell model. J Theor Biol. 2010;267: 605–13. doi: 10.1016/j.jtbi.2010.09.011. PMID:20837027

26. Ma W, Yu C, Zhang W, Zhou P, Hu J. The emergence of ribozymes synthesizing membrane components in RNA-based protocells. Biosystems. 2010;99: 201–9. doi: 10.1016/j.biosystems.2009.11.003. PMID:19961895

27. Bianconi G, Zhao K, Chen IA, Nowak MA. Selection for replicases in protocells. PLoS Comput Biol. 2013;9: e1003051. doi: 10.1371/journal.pcbi.1003051. PMID:23671413

28. Shah V, de Bouter J, Pauli Q, Tupper AS, Higgs PG. Survival of RNA Replicators is much Easier in Protocells than in Surface-Based, Spatial Systems. Life (Basel). 2019;9: 65. doi: 10.3390/life9030065. PMID:31394866

29. Johnson KA. A century of enzyme kinetic analysis, 1913 to 2013. FEBS Lett. 2013;587: 2753–66. doi: 10.1016/j.febslet.2013.07.012. PMID:23850893

30. Roussel MR. Heineken, Tsuchiya and Aris on the mathematical status of the pseudo-steady state hypothesis: A classic from volume 1 of Mathematical Biosciences. Math Biosci. 2019;318: 108274. doi: 10.1016/j.mbs.2019.108274. PMID:31697965

31. Conrad B, Iseli C, Pirovino M. Energy-harnessing problem solving of primordial life: Modeling the emergence of catalytic host-nested parasite life cycles. PloS One. 2023;18: e0281661. Doi: 10.1371/journal.pone.0281661. PMID:36972235

32. Briggs GE, Haldane JB. A Note on the Kinetics of Enzyme Action. Biochem J. 1925;19: 338–9. doi: 10.1042/bj0190338. PMID:16743508

33. Stubbs RT, Yadav M, Krishnamurthy R, Springsteen G. A plausible metal-free ancestral analogue of the Krebs cycle composed entirely of α-ketoacids. Nat Chem. 2020;12: 1016–1022. Doi: 10.1038/s41557-020-00560-7. PMID:33046840

34. Janzen E, Blanco C, Peng H, Kenchel J, Chen IA. Promiscuous Ribozymes and Their Proposed Role in Prebiotic Evolution. Chem Rev. 2020;120: 4879–4897. doi: 10.1021/acs.chemrev.9b00620. PMID:32011135

35. Tawfik DS. Accuracy-rate tradeoffs: how do enzymes meet demands of selectivity and catalytic efficiency? Curr Opin Chem Biol. 2014;21: 73–80. doi: 10.1016/j.cbpa.2014.05.008. PMID:24954689

36. Fitzsimmons WJ, Woods RJ, McCrone JT, Woodman A, Arnold JJ, Yennawar M, Evans R, Cameron CE, Lauring AS. A speed-fidelity trade-off determines the mutation rate and virulence of an RNA virus. PLoS Biol. 2018;16: e2006459. doi: 10.1371/journal.pbio.2006459. PMID:29953453

37. Furubayashi T, Ueda K, Bansho Y, Motooka D, Nakamura S, Mizuuchi R, Ichihashi N. Emergence and diversification of a host-parasite RNA ecosystem through Darwinian evolution. Elife. 2020;9: e56038. doi: 10.7554/eLife.56038. PMID:32690137

38. Mills DR, Peterson RL, Spiegelman S. An extracellular Darwinian experiment with a self-duplicating nucleic acid molecule. Proc Natl Acad Sci U S A. 1967;58: 217–24. doi: 10.1073/pnas.58.1.217. PMID:5231602

39. Marshall KA, Ellington AD. Molecular parasites that evolve longer genomes. J Mol Evol. 1999;49: 656–63. doi: 10.1007/pl00006586. PMID:10552046

40. Szilágyi A, Könnyű B, Czárán T. Dynamics and stability in prebiotic information integration: an RNA World model from first principles. Sci Rep. 2020;10: 51. doi: 10.1038/s41598-019-56986-8. PMID:31919467

41. Schreiber G, Haran G, Zhou HX. Fundamental aspects of protein-protein association kinetics. Chem Rev. 2009;109: 839–60. doi: 10.1021/cr800373w. PMID:19196002

42. Gleitsman KR, Sengupta RN, Herschlag D. Slow molecular recognition by RNA. RNA. 2017;23: 1745–1753. doi: 10.1261/rna.062026.117. PMID:28971853

43. Lawrence MS, Bartel DP. Processivity of ribozyme-catalyzed RNA polymerization. Biochemistry. 2003;42: 8748–55. doi: 10.1021/bi034228l. PMID:12873135

44. Martin LL, Unrau PJ, Müller UF. RNA synthesis by in vitro selected ribozymes for recreating an RNA world. Life (Basel). 2015;5: 247–68. doi: 10.3390/life5010247. PMID:25610978

45. Lukatsky DB, Shakhnovich BE, Mintseris J, Shakhnovich EI. Structural similarity enhances interaction propensity of proteins. J Mol Biol. 2007;365: 1596–606. doi: 10.1016/j.jmb.2006.11.020. PMID:17141268

46. McRae EKS, Wan CJK, Kristoffersen EL, Hansen K, Gianni E, Gallego I, Curran JF, Attwater J, Holliger P, Andersen ES. Cryo-EM structure and functional landscape of an RNA polymerase ribozyme. Proc Natl Acad Sci U S A. 2024;121: e2313332121. doi: 10.1073/pnas.2313332121. PMID:38207080

47. Qin S, Pang X, Zhou HX. Automated prediction of protein association rate constants. Structure. 2011;19: 1744–51. doi: 10.1016/j.str.2011.10.015. PMID:22153497

48. Koonin EV, Dolja VV, Krupovic M, Kuhn JH. Viruses Defined by the Position of the Virosphere within the Replicator Space. Microbiol Mol Biol Rev. 2021;85: e0019320. doi: 10.1128/MMBR.00193-20. PMID:34468181

49. de Sousa JAM, Fillol-Salom A, Penadés JR, Rocha EPC. Identification and characterization of thousands of bacteriophage satellites across bacteria. Nucleic Acids Res. 2023;51: 2759–2777. doi: 10.1093/nar/gkad123. PMID:36869669

50. Eppley JM, Biller SJ, Luo E, Burger A, DeLong EF. Marine viral particles reveal an expansive repertoire of phage-parasitizing mobile elements. Proc Natl Acad Sci U S A. 2022;119: e2212722119. doi: 10.1073/pnas.2212722119. PMID:36256808

51. Vignuzzi M, López CB. Defective viral genomes are key drivers of the virus-host interaction. Nat Microbiol. 2019;4: 1075–1087. doi: 10.1038/s41564-019-0465-y. PMID:31160826

52. Katzourakis A, Aswad A. The origins of giant viruses, virophages and their relatives in host genomes. BMC Biol. 2014;12: 51. doi: 10.1186/s12915-014-0051-y. PMID:25184667

53. Ibarra-Chávez R, Brady A, Chen J, Penadés JR, Haag AF. Phage-inducible chromosomal islands promote genetic variability by blocking phage reproduction and protecting transductants from phage lysis. PLoS Genet. 2022;18: e1010146. doi: 10.1371/journal.pgen.1010146. PMID:35344558

54. Rousset F, Depardieu F, Miele S, Dowding J, Laval AL, Lieberman E, Garry D, Rocha EPC, Bernheim A, Bikard D. Phages and their satellites encode hotspots of antiviral systems. Cell Host Microbe. 2022;30: 740-753.e5. doi: 10.1016/j.chom.2022.02.018. PMID:35316646

55. Genoyer E, López CB. The Impact of Defective Viruses on Infection and Immunity. Annu Rev Virol. 2019;6: 547–566. doi: 10.1146/annurev-virology-092818-015652. PMID:31082310

56. Fillol-Salom A, Miguel-Romero L, Marina A, Chen J, Penadés JR. Beyond the CRISPR-Cas safeguard: PICI-encoded innate immune systems protect bacteria from bacteriophage predation. Curr Opin Microbiol. 2020;56: 52–58. doi: 10.1016/j.mib.2020.06.002. PMID:32653777

57. Stauffer Y, Marguerat S, Meylan F, Ucla C, Sutkowski N, Huber B, Pelet T, Conrad B. Interferon-alpha-induced endogenous superantigen. a model linking environment and autoimmunity. Immunity. 2001;15: 591–601. doi: 10.1016/s1074-7613(01)00212-6. PMID:11672541

58. Malfavon-Borja R, Feschotte C. Fighting fire with fire: endogenous retrovirus envelopes as restriction factors. J Virol. 2015;89: 4047–50. doi: 10.1128/JVI.03653-14. PMID:25653437

59. Pirovino M, Iseli C, Curran J A, Conrad B. Biomathematical model of prebiotic autocatalytic RNA networks shows birth of adaptive immunity from degenerating molecular parasite catalysts. bioRxiv 023.05.25.542273; doi: 10.1101/2023.05.25.542273

60. Toti P, Sbordone L, Sbordone C, Bauer C. Ribozymes: analytical solution of the one-substrate, two-intermediate reversible scheme for enzyme reactions. J Biol Phys. 2006;32: 473–88. doi: 10.1007/s10867-006-9030-z. PMID:19669435

61. Morrison JF. Kinetics of the reversible inhibition of enzyme-catalysed reactions by tight-binding inhibitors. Biochim Biophys Acta. 1969;185: 269–86. doi: 10.1016/0005-2744(69)90420-3. PMID:4980133

62. Bar-Even A, Milo R, Noor E, Tawfik DS. The Moderately Efficient Enzyme: Futile Encounters and Enzyme Floppiness. Biochemistry. 2015;54: 4969–77. doi: 10.1021/acs.biochem.5b00621. PMID:26219075

