## Supporting Information for "Biomathematical enzyme kinetics model of prebiotic autocatalytic RNA networks: degenerating parasite-specific hyperparasite catalysts confer parasite resistance and herald the birth of molecular immunity"

**S1 Supporting Information**

**Related work**

Here we model an alternative temporal pathway leading up to parasite resistance according to which the hyperparasite preserves a small but significant catalytic activity. This model differs in several aspects with respect to the canonical proposition presented in the main text. Firstly, primary parasite resistance here is the interlinked consequence of the higher affinity binding of *F* to itself and *P* that at the same time reduces the catalytic activity according to the accuracy-rate trade-off [1]. Selectivity (accuracy, specificity) implies the ability to discriminate between two templates, provided both are present, and rate denotes speed [1-3]. Ground state discrimination means that specificity is achieved mainly through substrate binding, which imposes strong accuracy-rate trade-offs [1-2]. Improvements in selectivity mediated by tighter cognate substrate binding invariably lead to lower catalytic efficiency (parallel decreases in the constants for the cognate substrates *K_M_^cog^* and *k_cat_^cog^* [1]). This is precisely what the model stipulates, *F* has parasite- and hyperparasite-selectivity, which comes at the cost of a reduced catalytic activity that is expected, since it is a considerably smaller molecular species than the host molecules [4]. Secondly, while this solution offers superior parasite taming at the lower binding affinity level compared to the main model (compare A to C), by the very nature of the approach this comes at the expense of higher hyperparasite prevalence in the habitat (compare A to C); latter persists at low but significant levels even after binding affinities are one order of magnitude higher (compare B to D).

We add in Model 3 further terms for the catalytic activity of $F$:

$$F\Rightarrow\left( P\_F \right)\to^{c}F$$

$F\Rightarrow\left( F\_F \right)\to^{c}F$ $(3.0)$

Equation $\left( 3.4 \right),$ which governs the evolution of the hyperparasite $F=R_{4}$ becomes now:

$$\frac{dR_{4}}{dt}=(\beta k_{cat}^{1,3}R_{1}\_R_{3}+k_{cat}^{3,4}R_{3}\_R_{4}+k_{cat}^{4,4}R_{4}\_R_{4})\Pi-d_{R_{4}}R_{4} \left( 3.4' \right)$$

Here we choose$k_{cat}^{3,4}=k_{cat}^{4,4}=.04$with the same values of all other parameters (which can be freely chosen) as in Model 3.1 (Model 3.0.1 in Fig 1A in S1 Supporting Information).

Now, if the binding of the complexes $P\_F$, $F\_F$ is further strengthened, by e.g. raising$k_{on}^{3,4}=k_{on}^{4,4}=1.0$, and so also lowering the Michelis constants $K_{M}^{3,4}=K_{M}^{4,4}=0.14$, the situation for the host cycle can even be further stabilized, with a lower population of parasites $P$ and less molecules bound to the $P\_F$ and $F\_F$ complexes (Model 3.0.2 in Fig 1B in S1 Supporting Information). For comparison the main models without catalytic activity (Model 3.1 and 3.2 are shown in parallel in Figs 1C-D S1 Supporting Information.

**Supporting figures**

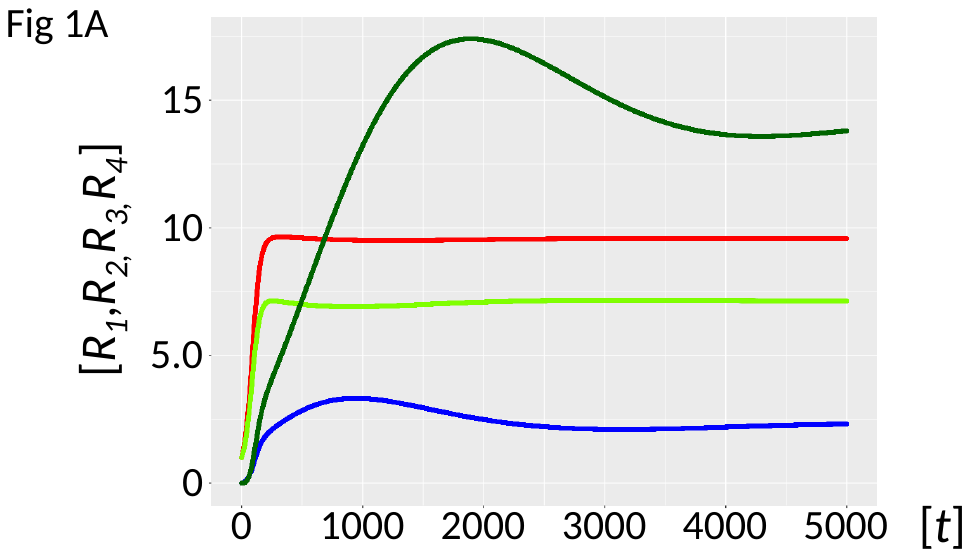

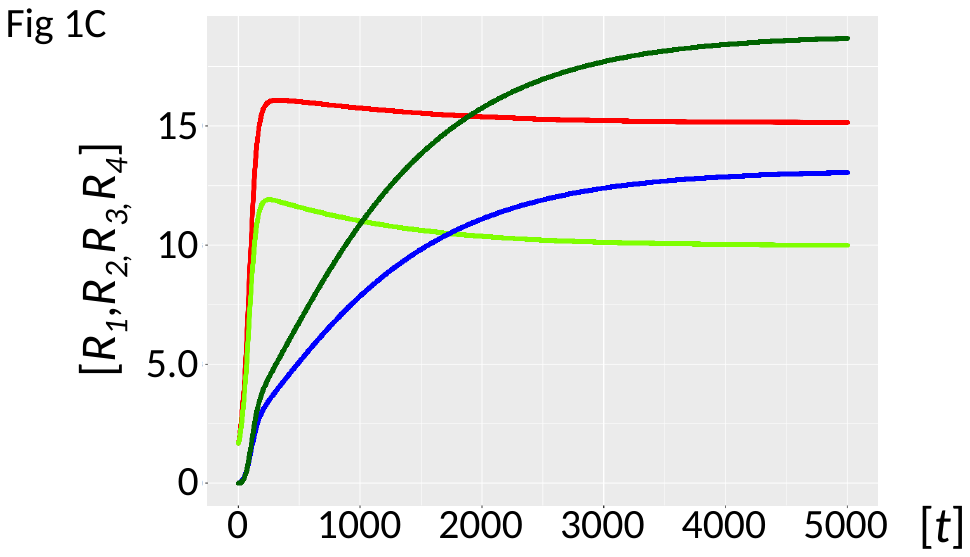

**Fig 1 in S1 Supporting Information.** Ribozyme concentrations of defined populations *R_1_*, *R_2_*, *R_3_* and *R_4_* [units on unspecified scale] are plotted against time *t* [units on unspecified scale]. Side by side comparison of parasite taming in two steps, A-B with reduced catalytic activity ($k_{cat}$ 0.04 sec^-1^), A with 1 μM *K_M_* values, and B with 0.14 μM. C-D denote the null catalytic values of the main model throughout ($k_{cat}$ 0.0 sec^-1^), and 1 μM *K_M_* values for C, and 0.14 μM for D. The ribozyme polymerase *R_1_* (red), the negative strand ribozyme *R_2_* (light green), the parasite *P* (blue), and the hyperparasite *F* (dark green).

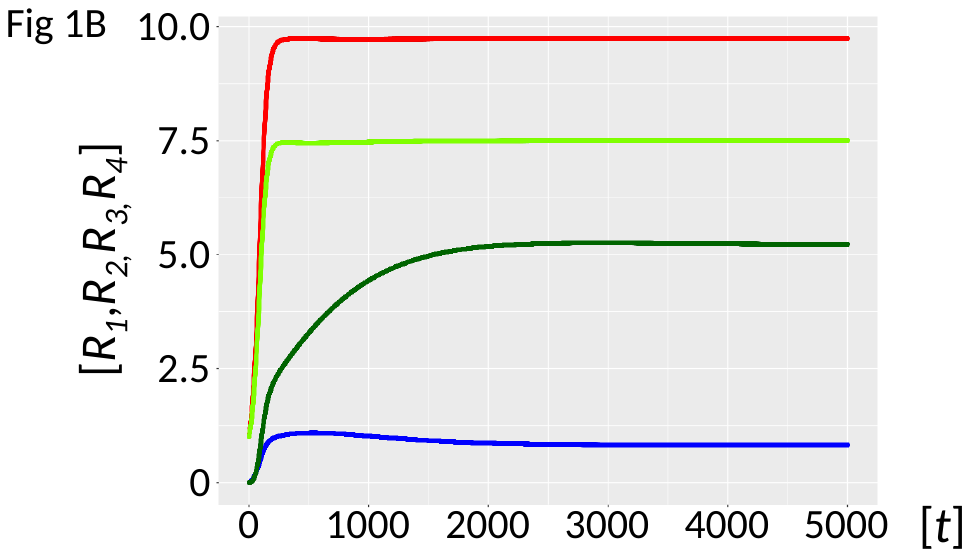

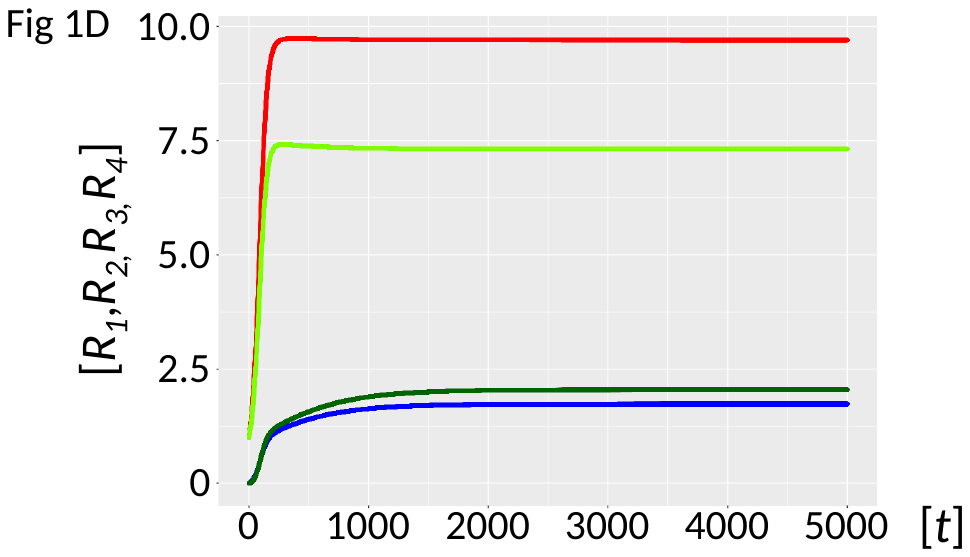

**Fig 1 in S1 Supporting Information.** Ribozyme concentrations of defined populations *R_1_*, *R_2_*, *R_3_* and *R_4_* [units on unspecified scale] are plotted against time *t* [units on unspecified scale]. Side by side comparison of parasite taming in two steps, A-B with reduced catalytic activity ($k_{cat}$ 0.04 sec^-1^), A with 1 μM *K_M_* values, and B with 0.14 μM. C-D denote the null catalytic values of the main model throughout ($k_{cat}$ 0.0 sec^-1^), and 1 μM *K_M_* values for C, and 0.14 μM for D. The ribozyme polymerase *R_1_* (red), the negative strand ribozyme *R_2_* (light green), the parasite *P* (blue), and the hyperparasite *F* (dark green).

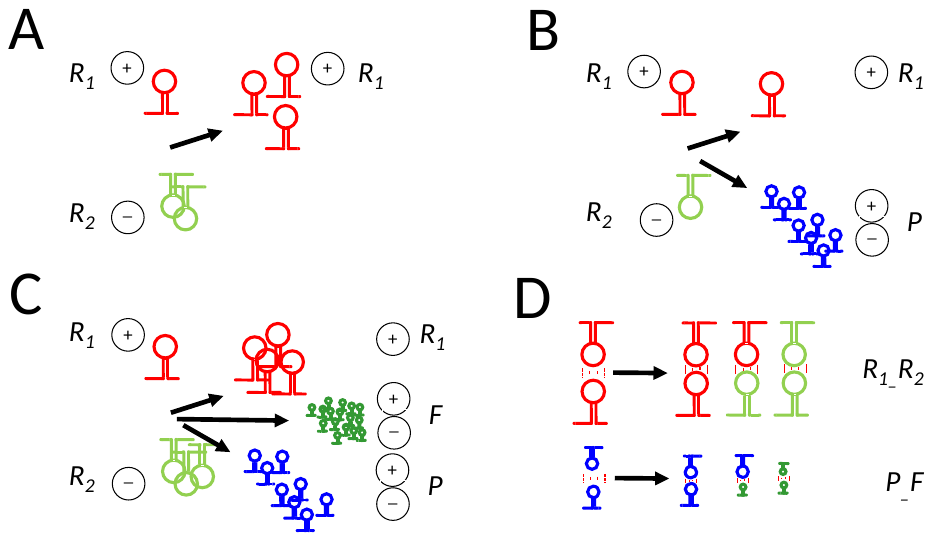

**Fig 2 in S1 Supporting Information.** The reactions schemes for the main model are graphically illustrated from A to D. **A is Model 1** showing the host cycle, provided sufficient building blocks are present. **B corresponds to Model 2** illustrating the emergence of molecular parasites *P*. **C is to Model 3** representing contemporaneous presence of hyperparasites *F* and parasites *P*. **D shows mechanistically how parasites are tamed** through increasing self-specific binding of host (upper row) and parasite/hyperparasites catalysts (lower row), from left (lower affinity) to right of both arrows (higher affinity). Interactions are depicted as kissing loop interactions [5].

**References**

1. Tawfik DS. Accuracy-rate tradeoffs: how do enzymes meet demands of selectivity and catalytic efficiency. Curr Opin Chem Biol. 2014;21: 73-80. doi: 10.1016/j.cbpa.2014.05.008. PMID:24954689
2. Janzen E, Blanco C, Peng H, Kenchel J, Chen IA. Promiscuous Ribozymes and Their Proposed Role in Prebiotic Evolution. Chem Rev. 2020;120: 4879-4897. doi: 10.1021/acs.chemrev.9b00620. PMID:32011135
3. Bar-Even A, Milo R, Noor E, Tawfik DS. The Moderately Efficient Enzyme: Futile Encounters and Enzyme Floppiness. Biochemistry. 2015;54: 4969-77. doi: 10.1021/acs.biochem.5b00621. PMID:26219075
4. Szilágyi A, Könnyű B, Czárán T. Dynamics and stability in prebiotic information integration: an RNA World model from first principles. Sci Rep. 2020;10: 51. doi: 10.1038/s41598-019-56986-8. PMID:31919467
5. McRae EKS, Wan CJK, Kristoffersen EL, Hansen K, Gianni E, Gallego I, Curran JF, Attwater J, Holliger P, Andersen ES. Cryo-EM structure and functional landscape of an RNA polymerase ribozyme. Proc Natl Acad Sci U S A. 2024;121: e2313332121. doi: 10.1073/pnas.2313332121. PMID:38207080
